# m^6^A Demethylase FTO Stabilizes LINK-A to Exert Oncogenic Roles via MCM3-Mediated Cell Cycle Progression and HIF-1α Activation

**DOI:** 10.1101/2023.01.28.526069

**Authors:** Yabing Nan, Shi Liu, Qingyu Luo, Xiaowei Wu, Pengfei Zhao, Wan Chang, Zhihua Liu

## Abstract

RNA *N*^6^-methyladenosine (m^6^A) modification, balanced by methyltransferases and demethylases, has recently been shown to play critical roles in multiple cancers. However, the mechanism by which m^6^A modification regulates long noncoding RNA (lncRNA) stability and function during cancer progression remains unclear. Here, we show that m^6^A demethylase fat mass and obesity-associated protein (FTO) removes the m^6^A modification on long intergenic noncoding RNA for kinase activation (LINK-A) and stabilizes it to promote cell proliferation and cytotoxic chemotherapy resistance in esophageal squamous cell carcinoma (ESCC). Mechanistically, LINK-A enhances the interaction between minichromosome maintenance complex component 3 (MCM3) and cyclin-dependent kinase 1 (CDK1) to promote MCM3 phosphorylation by CDK1. MCM3 is a subunit of the hexameric protein complex and its phosphorylation facilitates loading of the MCM complex onto chromatin, which promotes cell cycle progression and subsequent cell proliferation. Meanwhile, LINK-A prevents the interaction of MCM3 and hypoxia-inducible factor 1α (HIF-1α), abrogates MCM3-mediated transcriptional repression of HIF-1α, and promotes glycolysis and chemoresistance of cancer cells. These results elucidate a mechanism whereby FTO-stabilized LINK-A plays oncogenic roles and present the FTO/LINK-A/MCM3/HIF-1α axis as a promising therapeutic target for ESCC.

## Introduction

Esophageal cancer is a malignant tumor that arises from the esophagus, ranking seventh in incidence rate (approximately 604,000 new cases) and sixth in mortality (approximately 544,000 deaths) worldwide in 2020.^1^ There are two main histologic subtypes of esophageal cancer, esophageal squamous cell carcinoma (ESCC) and esophageal adenocarcinoma (EAC), with quite different etiologies.^2^ ESCC is the most prevalent histologic subtype, with poor prognosis and an extremely high prevalence worldwide.^2^ Chemotherapy is the first-line treatment for esophageal cancer, especially for patients with locally advanced cancer who have no indications for surgery; however, the benefits are unsatisfactory, and the five-year survival rate of patients with esophageal cancer is less than 20% due to chemoresistance.^2^ Thus, it is worth exploring novel oncogenic drivers and elucidating the potential molecular mechanisms to identify new therapeutic targets for ESCC.

*N*^6^-methyladenosine (m^6^A) modification is the most abundant internal RNA modification in eukaryotes, occurring mainly in messenger RNA (mRNA).^3,4^ Increasing evidence has shown that m^6^A modifications of noncoding RNAs (ncRNAs) such as microRNAs (miRNAs), long noncoding RNAs (lncRNAs) and circular RNAs (circRNAs) also play essential roles in various physiological and pathological bioprocesses including cancer development.^5-7^ m^6^A is installed by nuclear methyltransferases, termed “writers”, which are required for m^6^A formation.^4,8^ m^6^A-reader proteins such as YTH domain-containing proteins (YTHDCs) and insulin-like growth factor-2 mRNA-binding proteins (IGF2BPs) bind to m^6^A-containing RNAs to regulate their splicing, intracellular localization and stability.^9,10^ m^6^A methylation is dynamic and can be reversed by m^6^A demethylases, also named m^6^A “erasers”, including fat mass and obesity-associated protein (FTO) and α-ketoglutarate-dependent dioxygenase alkB homolog 5 (ALKBH5).^11,12^ Emerging evidence indicates that m^6^A modification exerts either oncogenic or tumor-suppressive effects under different conditions.^13,14^ However, the detailed mechanism by which m^6^A modification regulates ncRNAs during cancer progression remains largely elusive.

The switch from oxidative phosphorylation (OXPHOS) to aerobic glycolysis, called the Warburg effect, is the best characterized metabolic change in cancer cells.^15-17^ Although it is less efficient for ATP generation, aerobic glycolysis generates signaling metabolites to enhance cancer cell survival and therapeutic resistance under challenging conditions.^18,19^ Hypoxia inducible factor-1 (HIF-1), an oxygen-sensing transcription factor, determines whether glucose is consumed via oxidation or glycolysis.^20^ The oxygen-responsive HIF-1α subunit and constitutively expressed HIF-1β subunit form the heterodimeric HIF-1 transcription factor, which plays critical roles in the cellular response to hypoxia.^21^ The accumulation of HIF-1α and the activation of HIF-1α-mediated glucose transporters and rate-limiting enzymes in glucose metabolism reduce the efficiency of OXPHOS and promote glycolysis.^22-24^ Although it has been reported that transcriptional activation of HIF-1α leads to cancer metabolic reprogramming and therapeutic resistance,^25^ how lncRNAs regulate HIF-1α activation in ESCC remains unclear.

Here, we first identified that the m^6^A demethylase FTO demethylates and stabilizes a lncRNA long intergenic noncoding RNA for kinase activation (LINK-A) during ESCC chemoresistance. LINK-A directly interacts with minichromosome maintenance complex component 3 (MCM3) and cyclin-dependent kinase 1 (CDK1), which increases the phosphorylation of MCM3 at Ser112 by CDK1, and facilitates MCM3 incorporation into the MCM2-7 complex and subsequent chromatin loading, promoting cell cycle progression. On the other hand, LINK-A insulates the interaction between MCM3 and HIF-1α to free HIF-1α. The enhancement of HIF-1α transcriptional activity reprograms metabolic status from OXPHOS to glycolysis. Additionally, targeting LINK-A substantially sensitized ESCC patient-derived xenografts (PDXs) to first-line chemotherapy, thus indicating LINK-A is a promising target for cancer treatment.

## Results

### FTO stabilizes LINK-A and correlates with chemoresistance

To study the m^6^A modification of critical lncRNAs during ESCC progression, we first collected 34 baseline samples from ESCC patients who received neoadjuvant chemotherapy (17 were resistant and 17 were sensitive to chemotherapy) then detected the mRNA expression levels of two dominant m^6^A demethylases FTO and ALKBH5.^26^ The results indicated that FTO, but not ALKBH5, was significantly upregulated in chemoresistant samples compared with chemosensitive samples (Figures 1A and 1B). Moreover, we detected the protein level of FTO in the parental and chemoresistant ESCC cell lines previously established.^27^ The results showed that FTO protein expression was upregulated in both chemoresistant cell lines compared with their parental cells (Figure 1C). Furthermore, we detected m^6^A methylation using a m^6^A dot blot assay. As a result, the overall level of m^6^A was decreased in the chemoresistant cells compared with the parental cells (Figure 1D), which was consistent with the forced FTO expression in chemoresistant cells. Then, to further identified FTO-regulated lncRNAs, we depleted or overexpressed FTO in two ESCC cell lines and detected the expression levels of 14 annotated ncRNAs that were previously shown to be significantly upregulated in chemoresistant ESCC cell lines compared with the parental cell lines.^27^ The expression of four ncRNAs (TTLL3, LINC00663, PRSS30P and CETN4P) were too low to be detected by real-time quantitative PCR (RT‒qPCR). Among the remaining 10 ncRNAs, LINK-A was the only one found to be significantly downregulated following FTO knockdown and upregulated following FTO overexpression in both KYSE30 and KYSE450 cell lines (Figures 1E and 1F). Additionally, correlation analyses showed that LINK-A expression was positively correlated with FTO expression but barely correlated with ALKBH5 expression in the ESCC dataset (Figures S1A and S1B).

**Figure 1.**
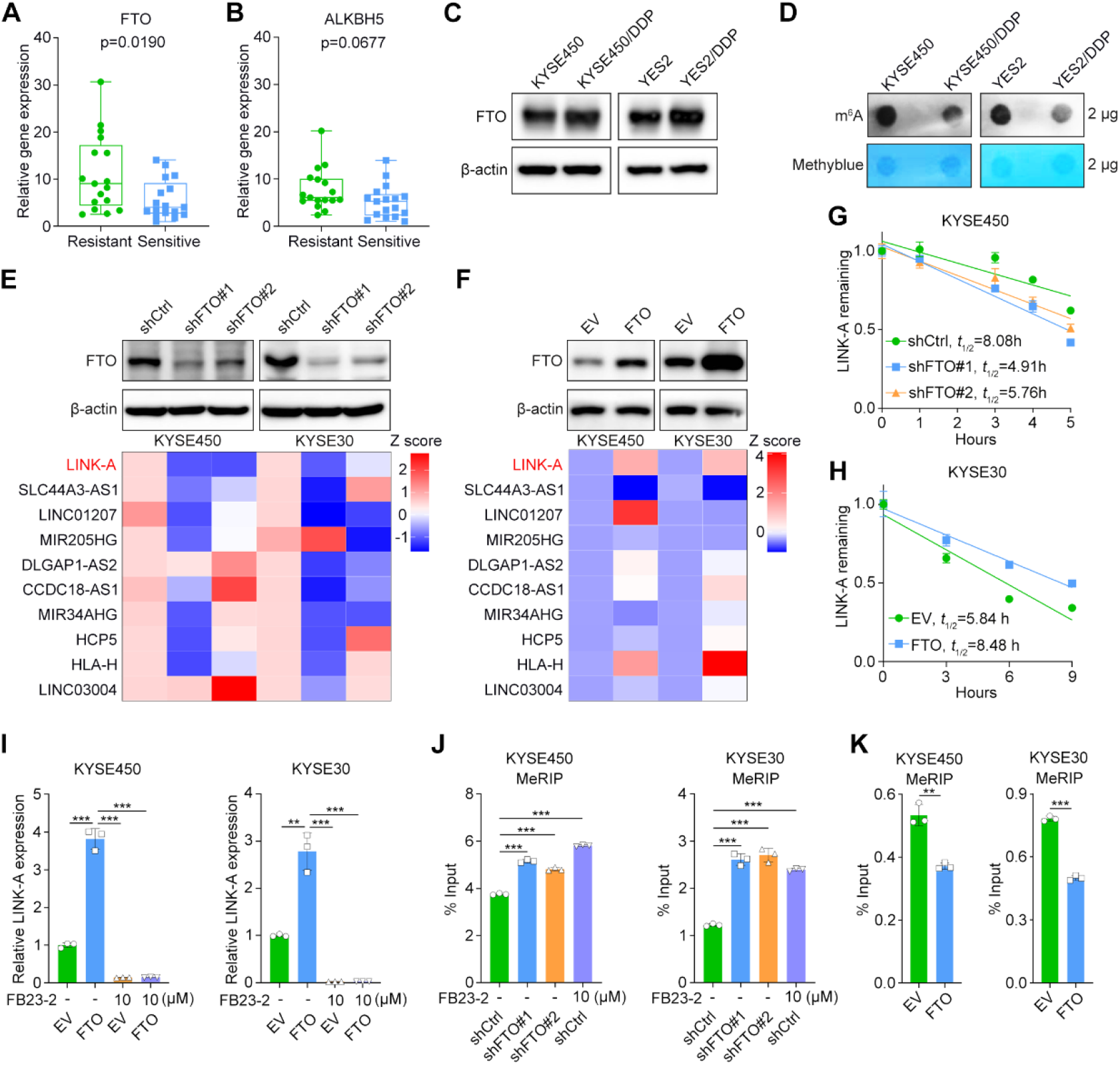
FTO stabilizes LINK-A and correlates with chemoresistance. (A and B) RT‒qPCR analysis of the expression of FTO (A) and ALKBH5 (B) in baseline samples from ESCC patients with chemosensitivity or chemoresistance. Box plot representation: from top to bottom—maximum, 75th percentile, median, 25th percentile and minimum values; unpaired Student’s t test; n = 17 for chemoresistant patients and n = 17 for chemosensitive patients. (C) Western blot analysis of FTO protein expression in chemoresistant and parental ESCC cell lines. (D) Total m^6^A level of chemoresistant and parental ESCC cell lines detected by m^6^A dot blot assay. Methylene blue staining was used as a loading control. (E and F) Western blot and heatmap of RT‒qPCR analyses in the indicated cells. (G and H) Kinetic analysis of RNA stability in the indicated cells. Actinomycin D (5 μg/ml) was added at time 0. The data are presented as the mean ± s.d. values; n = 3. (I) RT‒qPCR analysis of LINK-A expression in the indicated cells. Cells were pretreated with FB23-2 for 24 hours. The data are presented as the mean ± s.d. values; unpaired Student’s *t* test, ** p < 0.01, ***p < 0.001; n = 3. (J and K) MeRIP-qPCR analysis of LINK-A expression in the indicated cells. Control cells were pretreated with FB23-2 for 24 hours. The data are presented as the mean ± s.d. values; unpaired Student’s *t* test, ** p < 0.01, ***p < 0.001; n = 3.

To specify the mechanism by which FTO modulates LINK-A expression during ESCC chemoresistance, we first asked whether FTO affects LINK-A synthesis. No significant changes in LINK-A RNA synthesis were observed in FTO-depleted KYSE450 and KYSE30 cells (Figure S1C). Next, we performed an RNA stability assay and found that knockdown of FTO dramatically shortened the half-life of LINK-A and overexpression of FTO stabilized LINK-A (Figures 1G and 1H). Subsequently, we treated FTO-overexpressing KYSE30 and KYSE450 cells with FB23-2. The results indicated that overexpression of FTO markedly increased the expression level of LINK-A, whereas treatment with FB23-2 abrogated this effect (Figure 1I). At last, we determined whether the stabilization of LINK-A by FTO depended on its demethylase activity. MeRIP-qPCR results indicated that either knockdown of FTO or treatment with the FTO inhibitor FB23-2 significantly increased the m^6^A level of LINK-A, whereas overexpression of FTO decreased the m^6^A level of LINK-A (Figures 1J and 1K). These results indicated that FTO stabilized LINK-A expression in a m^6^A-dependent manner.

### LINK-A exerts oncogenic roles in ESCC

To investigated the roles of LINK-A in ESCC, we analyzed LINK-A expression in a cohort of ESCC patients by performing in situ hybridization (ISH). Survival analysis indicated that high LINK-A expression was dramatically correlated with poor prognosis in ESCC patients (Figure 2A). In addition, we found that LINK-A expression in cancer tissues was significantly higher than that in normal tissues (Figure 2B). However, there were no significant correlations of LINK-A expression with other clinicopathological features except for lymph node metastasis (Figure S2A). Moreover, the significant upregulation of LINK-A in cancer tissues was validated in the GEO database (Figure 2C). To further evaluate the potential of LINK-A as a therapeutic target in ESCC treatment, we first utilized an *in vitro* chemoresistant model. We determined the LINK-A expression level in long-term induced chemoresistant (KYSE450/DDP and YES2/DDP) and parental (KYSE450 and YES2) ESCC cell lines and found that the expression of LINK-A was dramatically upregulated 2- to 5-fold in the chemoresistant cells compared with the parental cells (Figure 2D). To determine the role of LINK-A in short-term drug exposure, we treated ESCC cells with DDP for different durations (0, 6, 12, 24 hours). The results indicated that LINK-A expression significantly increased as the DDP treatment time increased, especially after treatment for 24 hours (Figure 2E). These results suggested that LINK-A may exert crucial roles in both the acquisition and maintenance of chemoresistance.

**Figure 2.**
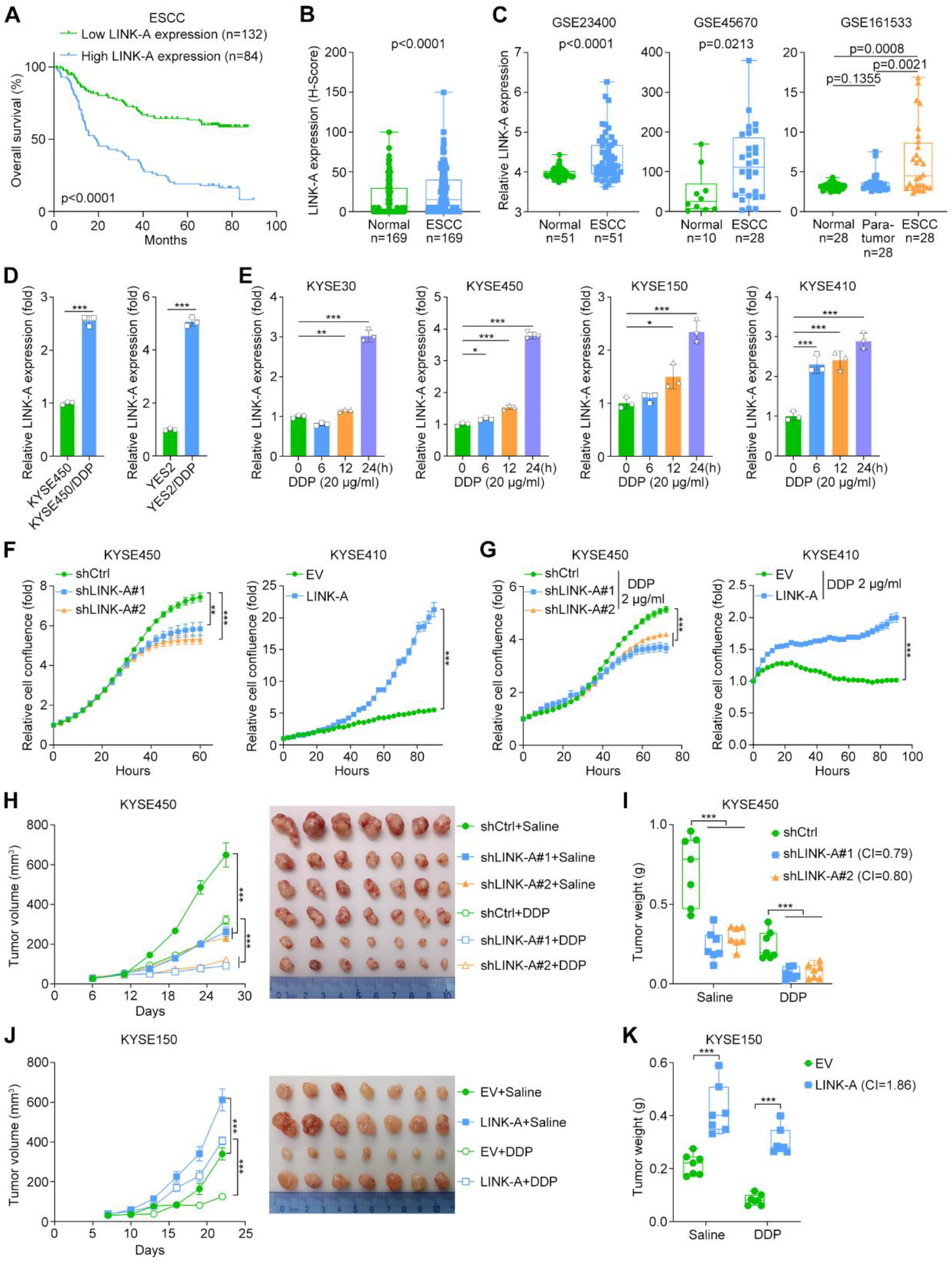
LINK-A exerts oncogenic roles in ESCC. (A) Overall survival of ESCC patients with low and high LINK-A expression levels (stratified by the mean expression level). Kaplan–Meier survival plots are shown. (B) Statistical analysis of LINK-A expression in 169 paired ESCC tissues and adjacent normal tissues. Box plot representation: from top to bottom—maximum, 75th percentile, median, 25th percentile and minimum values; paired Student’s *t* test. (C) Statistical analysis of LINK-A expression in ESCC tissues and adjacent normal tissues in the GEO database. Box plot representation: from top to bottom—maximum, 75th percentile, median, 25th percentile and minimum values; unpaired Student’s *t* test in GSE45670 and paired Student’s *t* test in GSE23400 and GSE161533. (D) RT–qPCR analysis of LINK-A expression in the indicated cells. The data are presented as the mean ± s.d. values; unpaired Student’s *t* test, ***p < 0.001; n = 3. (E) RT–qPCR analysis of LINK-A expression in the indicated cells treated with DDP for 0, 6, 12, and 24 hours. The data are presented as the mean ± s.d. values; unpaired Student’s *t* test, * p < 0.05, ** p < 0.01, ***p < 0.001; n = 3. (F and G) IncuCyte analysis of the indicated cells. Cells were treated with DDP at the indicated concentration in (G). The data are presented as the mean ± s.d. values; unpaired Student’s *t* test, ** p < 0.01, ***p < 0.001; n = 3. (H-K) Tumor growth curves and representative images (H and J) and tumor weights (I and K) of the xenografts derived from the indicated cells. The data are presented as the mean ± s.e.m. values in (H) and (J); Box plot representation: from top to bottom— maximum, 75th percentile, median, 25th percentile and minimum values in (I) and (K); unpaired Student’s *t* test, ***p < 0.001; n = 7.

Next, to verify the oncogenic roles of LINK-A, we performed *in vitro* and *in vivo* loss- and gain-of-function experiments. First, we established ESCC cell lines with stable LINK-A silencing and overexpression (Figures S2B and S2C). IncuCyte live cell image analysis showed that silencing of LINK-A significantly inhibited cell proliferation, whereas overexpression of LINK-A promoted cell proliferation (Figure 2F). Next, to evaluate the effect of LINK-A on chemoresistance, we treated ESCC cells with cisplatin (DDP) and found that LINK-A significantly increased resistance to cytotoxic chemotherapy (Figure 2G). Finally, xenograft assays indicated that silencing LINK-A dramatically retarded tumor growth and elicited chemosensitivity, while overexpression of LINK-A significantly promoted proliferation and resistance to cytotoxic chemotherapy (Figures 2H-2K). Taken together, these results indicated that LINK-A plays oncogenic roles in ESCC.

### LINK-A facilitates MCM3 phosphorylation by enhancing the interaction between MCM3 and CDK1

To further illustrate the downstream regulatory mechanism by which LINK-A plays an oncogenic role, we performed MS analyses to identify LINK-A-precipitated proteins in KYSE450 and YES2 cells. In total, 43 and 14 LINK-A-interacting proteins were identified in KYSE450 and YES2 cells, respectively, and 7 of them were consistently identified in both cell lines (Figure 3A). Among these 7 proteins, the MCM3 has been reported to play an important role in the cell cycle regulation and DNA replication,^28^ whereas its detailed molecular mechanisms in ESCC, especially in mediating cancer cell proliferation and chemoresistance, remain elusive. Next, we performed RNA pulldown followed by Western blotting to confirm the specific interaction between MCM3 and LINK-A (Figure 3B). Moreover, this interaction was validated in KYSE30 and KYSE450 cells by RIP-qPCR (Figure 3C). Fluorescence in situ hybridization (FISH) and immunofluorescence (IF) double staining showed that endogenous LINK-A and MCM3 were colocalized in the nucleus (Figure S3A). Next, we explored the effect of LINK-A on its interacting protein MCM3. Knockdown or overexpression of LINK-A did not affect either the RNA level or protein level of MCM3 (Figures 3D, 3E and S3B-S3E). Strikingly, Western blot analysis indicated that LINK-A markedly facilitated MCM3 phosphorylation at Ser112 (Figures 3D and 3E). Moreover, the phosphorylation level of MCM3 at Ser112 was increased in a dose-dependent manner upon transient LINK-A overexpression (Figures 3F and 3G).

**Figure 3.**
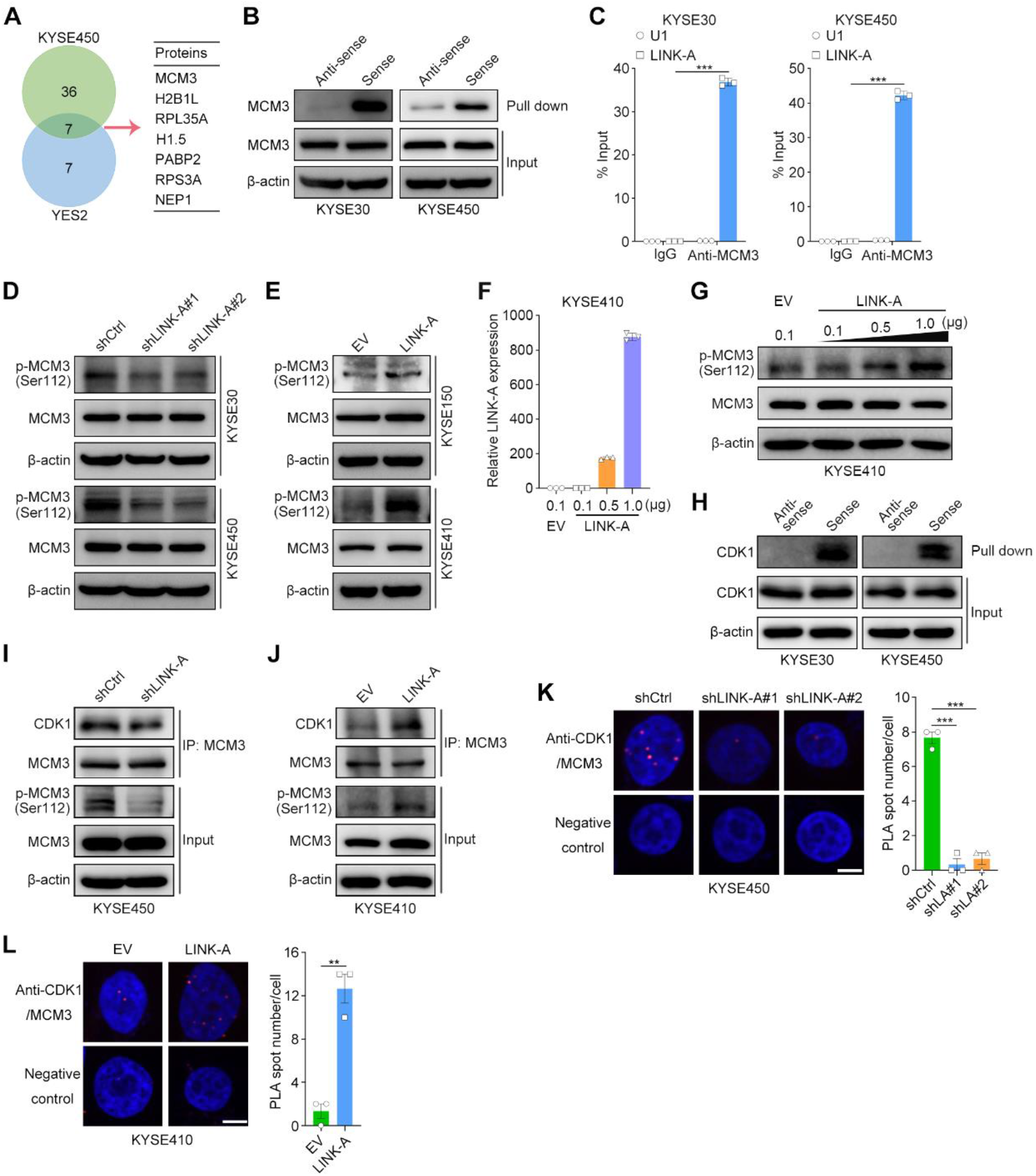
LINK-A promotes MCM3 phosphorylation by enhancing the interaction between MCM3 and CDK1. (A) A list of LINK-A-associated proteins identified by MS following RNA pulldown assays in KYSE450 and YES2 cells. (B) Western blot analysis of MCM3 expression following RNA pulldown assays in the indicated cells. (C) RT‒qPCR analysis of LINK-A and U1 expression following RIP assays in the indicated cells. The data are presented as the mean ± s.d. values; unpaired Student’s *t* test, ***p < 0.001; n = 3. (D and E) Western blot analysis of MCM3 and phosphorylated MCM3 in the indicated cells. (F and G) RT‒qPCR analysis of LINK-A expression (F) and Western blot analysis of MCM3 and phosphorylated MCM3 (G) after exogenous expression of increasing amounts of LINK-A in KYSE410 cells. (H) Western blot analysis of CDK1 expression following RNA pulldown assay in the indicated cells. (I and J) Western blot analysis of IP results in the indicated cells. (K and L) Representative images of the merged PLA and nuclear (DAPI) channels and quantitative analysis of PLA spots in the indicated cells. The scale bar represents 10 μm. The data are presented as the mean ± s.e.m. values; unpaired Student’s *t* test, ** p < 0.01, ***p < 0.001; n = 3.

A previous study reported that CDK1 phosphorylates MCM3 at Ser112 and mediates MCM2-7 complex assembly and chromatin loading.^29^ We then determined whether LINK-A promotes MCM3 phosphorylation by acting as a scaffold for CDK1 and MCM3. The results of RNA pulldown followed by Western blotting confirmed the interaction between LINK-A and CDK1 in KYSE30 and KYSE450 cells (Figure 3H). In addition, FISH and IF double staining indicated colocalization of endogenous LINK-A and CDK1 in the nucleus (Figure S3F). Next, the co-IP assay indicated that knockdown of LINK-A dramatically suppressed and that overexpression of LINK-A significantly increased the interaction between CDK1 and MCM3 (Figures 3I and 3J). Furthermore, proximity ligation assay (PLA) confirmed that LINK-A promotes the interaction between CDK1 and MCM3 (Figures 3K and 3L).

### Phosphorylated MCM3 mediates the oncogenic roles of LINK-A

To determine whether LINK-A-mediated MCM3 phosphorylation promotes MCM3 complex assembly and subsequent chromatin loading in ESCC, we performed cellular fractionation and then detected chromatin-bound proteins by Western blot analysis. The results showed that knockdown of LINK-A dramatically decreased the chromatin loading of MCM2, MCM3, MCM4, MCM5, MCM6, and MCM7 proteins, whereas overexpression of LINK-A promoted chromatin loading of the MCM2-7 complex in ESCC cells (Figures 4A and 4B). Next, we constructed a Ser112 site mutant of MCM3 and transfected MCM3 wild-type MCM3 and MCM3^S112A^ (Ser112 mutated to alanine) into endogenous MCM3-depleted cells. The increased chromatin loading of the MCM2-7 complex mediated by LINK-A was abolished by mutation at the Ser112 site, indicating this process is dependent on MCM3 phosphorylation (Figure 4C). To specify whether MCM3 mediates the cell cycle progression driven by LINK-A, we performed rescue assays and the results showed that overexpression of MCM3, but not MCM3^S112A^, completely abolished LINK-A silencing-mediated G0/G1 arrest (Figures 4D and S4A, S4B). In contrast, silencing MCM3 in LINK-A-overexpressing KYSE410 cells rescued LINK-A-mediated cell cycle progression (Figures 4E and S4C, S4D). Furthermore, IncuCyte live cell image analysis indicated that wild-type MCM3, but not its mutant, completely mediated the ability of LINK-A to promote the proliferation of KYSE450 and KYSE410 cells (Figures 4F and 4G). To further demonstrate the functional relationship between MCM3 and LINK-A in vivo, we performed xenograft assays with KYSE150 cells (Figure S4E). The results indicated that overexpression of LINK-A markedly promoted cell proliferation and chemoresistance, whereas silencing of MCM3 abrogated the promoting effects of LINK-A on cell proliferation and chemoresistance (Figures 4H and 4I). Taken together, these results indicated that LINK-A exerted oncogenic effects by enhancing MCM3 phosphorylation to facilitate MCM2-7 complex assembly and subsequent chromatin loading.

**Figure 4.**
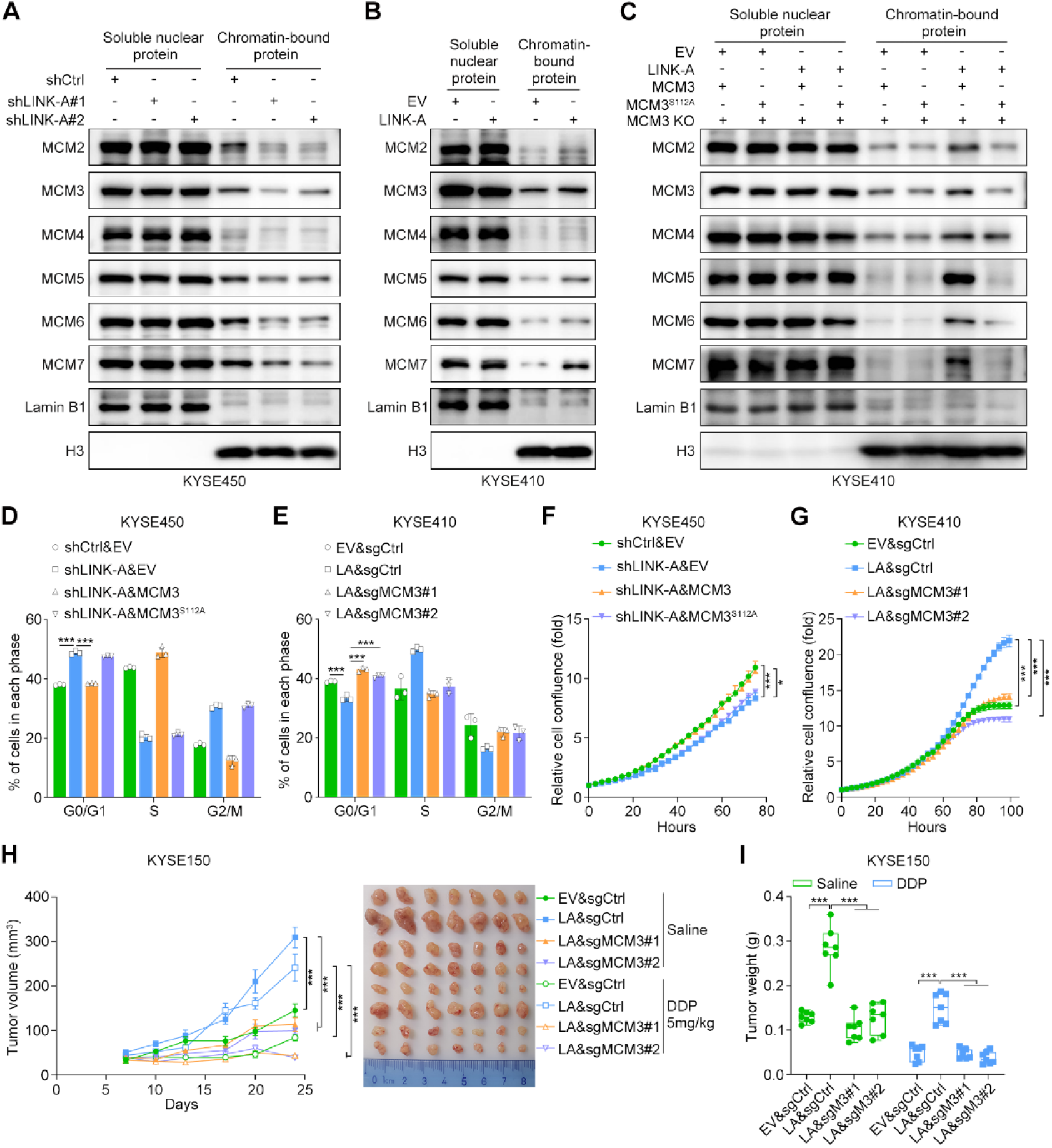
Phosphorylated MCM3 mediates the oncogenic roles of LINK-A. (A-C) Western blot analysis of the nuclear and chromatin-bound proteins in the indicated cells. (D and E) Quantitative analysis of the cell cycle distribution (percentage) in the indicated cells. The data are presented as the mean ± s.d. values; unpaired Student’s *t* test, ***p < 0.001; n = 3. (F and G) IncuCyte analysis of the indicated cells. The data are presented as the mean ± s.d. values; unpaired Student’s *t* test, * p < 0.05, ***p < 0.001; n = 3. (H and I) Tumor volumes and representative images (H) and tumor weights (I) of the xenografts derived from the indicated cells. The data are presented as the mean ± s.e.m. values in (H); Box plot representation: from top to bottom—maximum, 75th percentile, median, 25th percentile and minimum values in (I); unpaired Student’s *t* test, ***p < 0.001; n = 7.

### LINK-A induces HIF-1α transcriptional activity by sequestering HIF-1α from MCM3

To further study the mechanism how LINK-A/MCM3 leads to the resistance to cytotoxic chemotherapy, we performed RNA-seq in LINK-A-silenced KYSE450 cells. Strikingly, the gene set enrichment analysis (GSEA) showed that glycolysis was significantly enriched in control cells compared with two knockdown arms mediated by two independent shRNA sequences (Figure 5A). Given the observation in a previous study that MCM3 negatively regulates HIF-1α transcriptional activity by directly interacting with HIF-1α,^30^ we asked whether LINK-A activates HIF-1α to promote tumor glycolysis. First, we determined whether LINK-A influences the interaction between MCM3 and HIF-1α by performing IP assays in LINK-A-silenced KYSE450 cells and LINK-A-overexpressing KYSE410 cells. The results suggested that silencing LINK-A markedly increased while overexpression of LINK-A suppressed the interaction between MCM3 and HIF-1α (Figure 5B). Moreover, we purified MCM3 and HIF-1α proteins by GST pulldown assay and demonstrated that the LINK-A sense strand but not the antisense strand suppressed the interaction between MCM3 and HIF-1α in vitro (Figure 5C). Since the expression of HIF-1α was not altered by either overexpression or knockdown of LINK-A, we hypothesized that LINK-A may regulate HIF-1α by activating its transcriptional activity (Figures S5A and S5B). Accordingly, silencing of LINK-A markedly suppressed and overexpression of LINK-A enhanced hypoxia-responsive element (HRE)-luciferase activity (Figures 5D and 5E). Moreover, RT‒qPCR results indicated that LINK-A dramatically facilitated the expression of HIF-1α target genes, including SLC2A1, PDK1, BCL2, HK2, LDHA and VEGF (Figures 5F and 5G).

**Figure 5.**
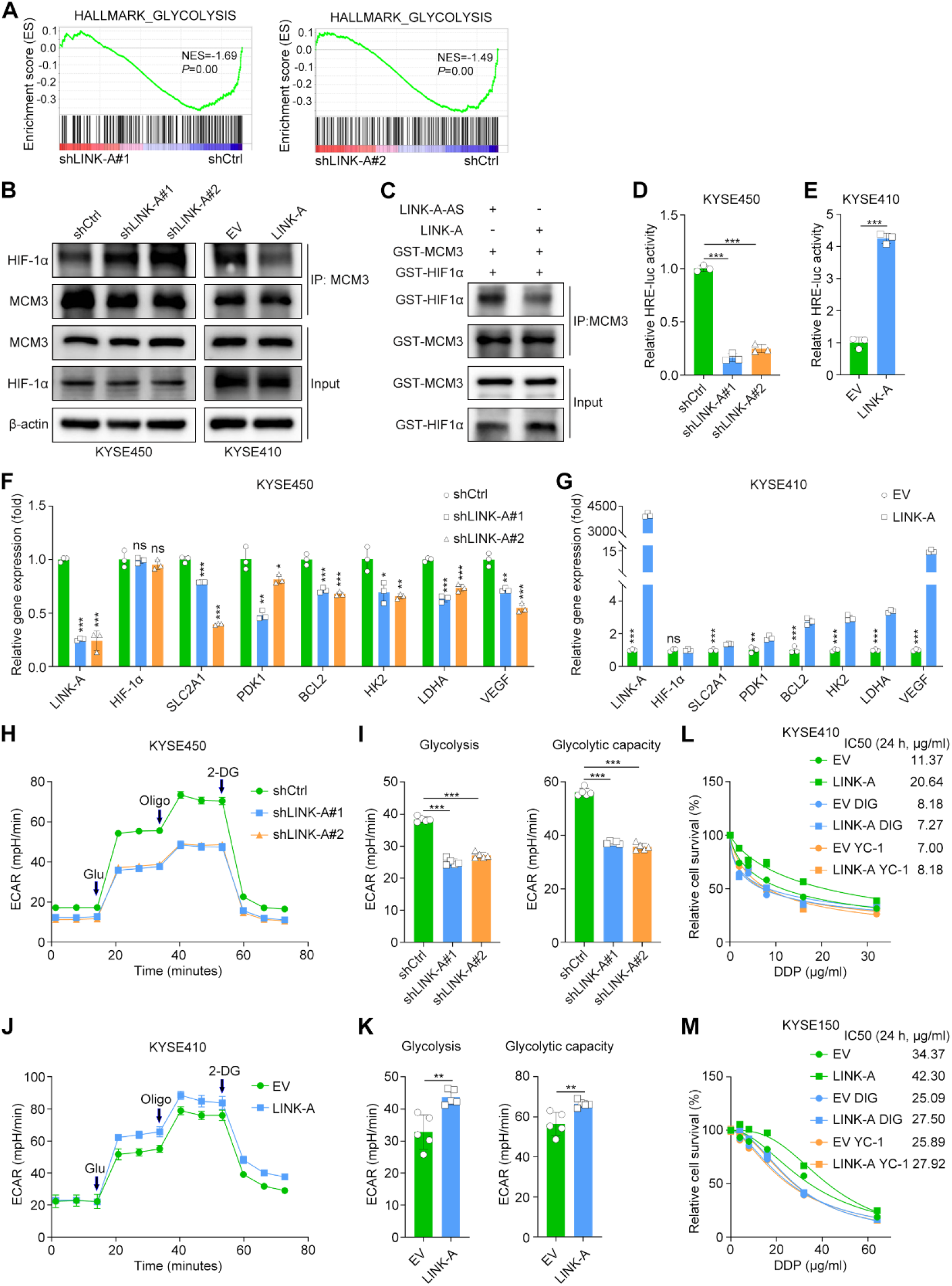
LINK-A induces HIF-1α transcriptional activity by sequestering HIF-1α from MCM3. (A) Pathways enriched by GSEA based on RNA-seq data from LINK-A-silenced KYSE450 cells. (B and C) Western blot analysis of the interaction of MCM3 and HIF-1α in vivo (B) and in vitro (C). (D and E) Relative HRE-luciferase activity in the indicated cells. The data are presented as the mean ± s.d. values; unpaired Student’s *t* test, ***p < 0.001; n = 3. (F and G) RT‒qPCR analyses of HIF-1α target genes in the indicated cells. The data are presented as the mean ± s.d. values; unpaired Student’s *t* test, * p < 0.05, ** p < 0.01, ***p < 0.001, ns = no significant difference; n = 3. (H-K) Measurement of the ECAR and glycolysis levels using a Seahorse assay in LINK-A-silenced KYSE450 cells (H and I) and LINK-A-overexpressing KYSE410 cells (J and K). The data are presented as the mean ± s.d. values; unpaired Student’s *t* test, ** p < 0.01, ***p < 0.001; n = 5. (L and M) Relative viability of the indicated cells after DDP treatment for 24 hours. Cells were pretreated with lificiguat (YC-1, 100 μM) or digoxin (100 nM) for 48 hours. The data are presented as the mean ± s.d. values; n = 3.

Given the important roles of HIF-1α in cancer metabolic reprogramming, we performed a Seahorse assay to determine the effect of LINK-A on metabolic reprogramming in ESCC cells. The results showed that knockdown of LINK-A significantly decreased the extracellular acidification rate (ECAR) but moderately increased the oxygen consumption rate (OCR) (Figures 5H, 5I and S5C, S5D). Similarly, overexpression of LINK-A increased the ECAR but decreased the OCR (Figures 5J, 5K and S5E, S5F). Lastly, we examined whether HIF-1α mediates the role of LINK-A in chemoresistance in ESCC. The cell viability assay showed that treatment with the HIF-1α inhibitors lificiguat (YC-1) and digoxin markedly suppressed the oncogenic effects of LINK-A in KYSE410 and KYSE150 cells (Figures 5L and 5M). Taken together, these results suggested that LINK-A activated HIF-1α by abrogating the MCM3-mediated transcriptional repression of HIF-1α and then facilitated glycolysis and chemoresistance.

### Targeting LINK-A sensitizes ESCC to cytotoxic chemotherapy

To determine whether targeting LINK-A is a promising treatment strategy for ESCC, we established three PDX models. We found that adeno-associated virus (AAV)-mediated silencing of LINK-A dramatically increased DDP sensitivity in ESCC (Figures 6A-6F). Finally, we detected the RNA expression of LINK-A in the same baseline samples as showed in Figure 1A. The result indicated that the expression level of LINK-A was significantly upregulated in the chemoresistant samples compared with chemosensitive samples (Figure 6G). Furthermore, correlation analysis showed that the expression of LINK-A was positively correlated with the expression of FTO in these samples (Figure 6H), which further confirmed the epigenetic regulation of FTO on LINK-A during ESCC chemoresistance. Taken together, our study identified LINK-A as a chief culprit for ESCC chemoresistance and revealed the potential mechanism of LINK-A-mediated chemoresistance and epigenetic regulation of LINK-A in ESCC (Figure 6I).

**Figure 6.**
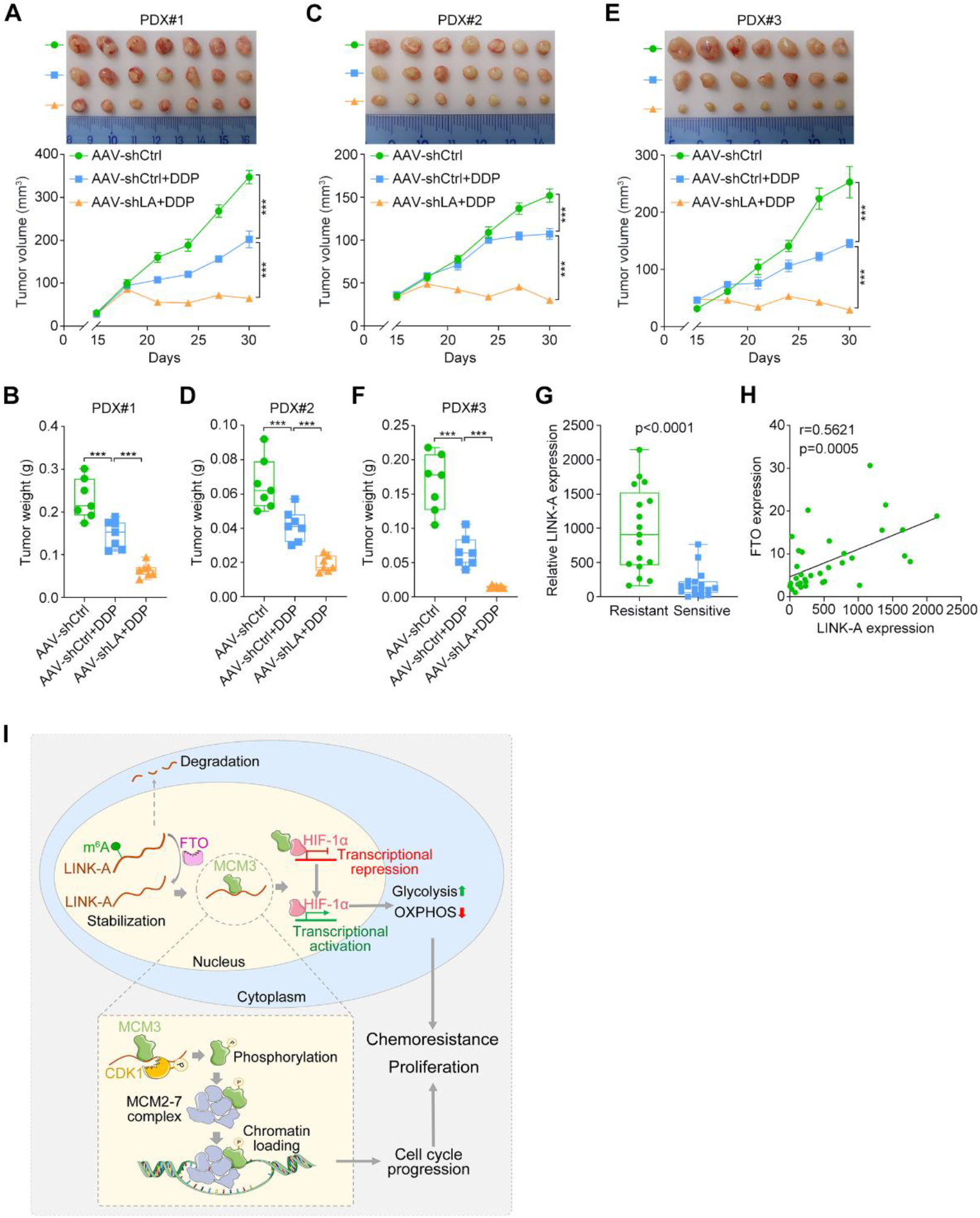
Targeting LINK-A sensitizes ESCC to cytotoxic chemotherapy. (A-F) Tumor growth curves and representative images (A, C and E) and tumor weights (B, D and F) in three PDX models of ESCC. AAVs were intratumorally injected every three days for a total of two treatments. The data are presented as the mean ± s.e.m. values in (A), (C) and (E); Box plot representation: from top to bottom—maximum, 75th percentile, median, 25th percentile and minimum values in (B), (D) and (F); unpaired Student’s t test, ***p < 0.001; n = 7. (G) RT‒qPCR analysis of the expression of LINK-A in baseline samples from ESCC patients with chemosensitivity or chemoresistance. Box plots representation: from top to bottom—maximum, 75th percentile, median, 25th percentile and minimum values; unpaired Student’s t test; n = 17 for chemoresistant patients and n = 17 for chemosensitive patients. (H) Correlation analysis between the expression of the FTO and LINK-A in ESCC samples. Spearman correlation coefficients are shown. (I) Schematic summary of the mechanism by which the FTO-LINK-A-MCM3-HIF-1α axis mediates ESCC malignancy.

## Discussion

The m^6^A demethylase FTO has been reported to be a tumor-promoting factor in numerous cancers,^31-33^ whereas ALKBH5 has been identified to exert either oncogenic or tumor-suppressive roles.^34-36^ In the current study, we identified FTO as a significantly upregulated m^6^A demethylase in chemoresistant ESCC patients compared with chemosensitive patients. Integrating our previous data that 14 annotated ncRNAs were significantly upregulated in chemoresistant ESCC cell lines compared with the parental cell lines,^27^ we identified LINK-A as the most significantly stabilized lncRNA by FTO. As an oncogenic lncRNA, LINK-A has been reported to activate diverse cancer-related pathways in several cancers.^37-40^ Although the oncogenic roles of LINK-A have been proposed, the epigenetic regulation of LINK-A, including m^6^A methylation, has not yet been reported. Herein, we found that FTO stabilizes LINK-A expression via m^6^A demethylation and elucidated the oncogenic roles of LINK-A that contribute to cancer cell proliferation and chemoresistance in ESCC.

LINK-A activates HIF-1α signaling and promotes glycolytic reprogramming and tumorigenesis by facilitating the activation of breast tumor kinase in TNBC.^37^ In addition, LINK-A can activate HIF-1α in osteoasrcoma^41^ and diabetic nephropathy.^42^ Conversely, the MCM3 protein directly binds to the HIF-1α subunit and synergistically inhibits HIF-1α transcriptional activity via distinct O_2_-dependent mechanisms.^30^ Here, we identified a novel mechanism by which LINK-A indirectly activates HIF-1α by interacting with MCM3 and insulating the MCM3-mediated transcriptional repression of HIF-1α. Transcriptional activation of HIF-1α contributes to metabolic reprogramming from OXPHOS to glycolysis in cancer cells and eventually promotes chemoresistance. Interesting, we found that the interaction between LINK-A and MCM3 leads to cell cycle progression and subsequent cell proliferation via another independent mechanism. MCM3 is a member of the MCM family, which was originally identified through a yeast genetic screen.^43^ The MCM2-7 proteins are highly conserved and form a hexameric complex during the G1 phase; this complex is then loaded onto the origin of replication to form a prereplication complex that is indispensable for the initiation of DNA replication.^44^ CDK1-dependent MCM3 phosphorylation at Ser112 triggers the association of MCM3 with the remaining MCM subunits and subsequent chromatin loading of the MCM2-7 complex.^29^ Our study indicated that LINK-A acts as a protein‒protein scaffold to mediate CDK1-dependent phosphorylation of MCM3 at Ser112 and increase the chromatin loading of MCM3, which in turn promotes cell cycle progression and cell proliferation.

In conclusion, this study reveals the details regarding m^6^A demethylation modification of LINK-A in ESCC. The m^6^A demethylase FTO mediates the stabilization and supports the oncogenic roles of LINK-A. The direct interaction between LINK-A and MCM3 not only increases CDK1-mediated phosphorylation of MCM3 at Ser112, which promotes the assembly and chromatin loading of the MCM2-7 complex and subsequent cell cycle progression and cell proliferation, but also insulates MCM3-mediated HIF-1α transcriptional repression, which facilitates tumor metabolic reprogramming and chemoresistance. Collectively, our findings suggest that the FTO/LINK-A/MCM3/HIF-1α axis is crucial for the progression of ESCC, and this discovery provides a promising therapeutic strategy for targeting LINK-A in ESCC patients, especially those with high LINK-A expression levels.

### Materials and Methods Patient samples and ISH

A total of 34 baseline samples from patients treated with neoadjuvant chemotherapy were collected from the Cancer Hospital, Chinese Academy of Medical Sciences. Written informed consent was obtained from all patients. Human ESCC tissue microarray slides were purchased from Servicebio (#G6040; Wuhan, Hubei, China). RNA ISH was performed as previously described.^27^ The sequences of the probes used for ISH were as follows (5’ to 3’): Probe #1, CACTAGGGTGGAACCTCAGGAAGTTGATGACATTTGTAGC; Probe #2, GCTTATAATTCTTCTAATTACTAACAATGCTGGTATGAAA; and Probe #3, GACTCGCTCTGGCCGTATGTAATGATGTCTGTGGCTACAT.

### Cell culture

The human ESCC cell lines were kindly provided by Dr. Yutaka Shimada (Kyoto University, Kyoto, Japan). The 293T cell line was purchased from the American Type Culture Collection (ATCC; Manassas, VA, USA). ESCC cells were cultured in RPMI-1640 medium supplemented with 10% fetal bovine serum (FBS), and 293 T cells were cultured in Dulbecco’s modified Eagle’s medium (DMEM) supplemented with 10% FBS. All cells were cultured under aseptic conditions at 37 °C in 5% CO_2_ and passaged with 0.25% trypsin. All cell lines were routinely verified using short tandem repeat DNA fingerprinting and were tested for *Mycoplasma* contamination using a MycoBlue Mycoplasma Detector (D101; Vazyme Biotech, Nanjing, Jiangsu, China) before use in any experiments.

### Lentivirus production and infection

The full-length cDNA of LINK-A was synthesized using PCR amplification and subsequently inserted into the pLVX-IRES-neo vector (#632184; Clontech, Mountain View, CA, USA). The full-length cDNA of MCM3 was cloned into the plasmid PEZ-LV105 (M0227; GeneCopoeia, Guangzhou, Guangdong, China). The shRNA sequences were cloned into the pSIH1-puro vector (#26597; Addgene, Cambridge, MA, USA). The sgRNA sequences were cloned into the LentiCRISPR v.2 (#52961; Addgene) backbone. For the construction of the MCM3^S112A^ mutant, the plasmid containing the MCM3 OFR was used as the template for PCR. The first round of PCR used two pairs of primers containing the mutation site, and the second round of PCR used primers at both ends. Finally, the obtained fragment was cloned into the vector by restriction endonuclease ligation and the mutation site was verified by Sanger sequencing. The sequences of the PCR primers used for mutant construction were as follows (5’ to 3’): round#1-F: CGGAATTCGCCACCATGGCGGGTACCGTGGTGCT; round#1-R: GCTCTAGATCATTGTCATCGTCGTCCTTGTAATCCTAGATGAGGAAGATGATGC; round#2-F: AAGCACGTCGCCCCGCGGAC; round#2-R: GTCCGCGGGGCGACGTGCTT. Transfection was performed using Hieff Tranns Liposomal Transfection Reagent (40802, Yeasen, Shanghai, China) according to the manufacturer’s protocol. Lentiviruses were produced in 293T cells with a second-generation packaging system containing psPAX2 (#12260; Addgene) and pMD2.G (#12259; Addgene). Cells were infected with lentivirus in the presence of 8 μg/ml polybrene (Sigma–Aldrich, St. Louis, MO, USA) twice within a 48-hour period and were then selected with 1 μg/ml puromycin (Sangon Biotech, Shanghai, China) or 200 μg/ml G418 (Sigma–Aldrich) for 7 days. The shRNA and sgRNA sequences are listed in Supplementary Table S1.

### Cell viability assay

The cell proliferation rate and cell survival were assessed as previously described.^27^ For the inhibitor (YC-1 (HY-14927; MedChemExpress) and digoxin (HY-B1049; MedChemExpress)) treatment experiments, cells were treated with the inhibitors at the corresponding concentrations for 48 hours before seeding into 96-well plates. Cell Counting Kit-8 reagent (CK04; Dojindo Laboratories, Kumamoto, Japan) was used to assess the cell proliferation rate and cell viability after DDP treatment according to the instructions.

### Live cell analysis

An IncuCyte Live Cell Imaging System (Essen Bioscience, Hertfordshire, UK) was used for live cell imaging. Cells were seeded in 96-well plates, and drugs were added after cell adhesion. The plates were placed in the IncuCyte system, and snapshots of two regions per well were acquired at 3-hour intervals over a 72-hour period. Cell confluence was quantified with the accompanying commercial software.

### Cell cycle analysis

Cell cycle analysis was performed using a Cell Cycle Assay Kit (C543; Dojindo Laboratories) according to the manufacturer’s instructions. The cell cycle was analyzed using a flow cytometer.

### Western blot and RT–qPCR analyses

Western blot analysis was performed as previously described.^45^ In brief, ESCC cells were lysed with RIPA buffer (CW2333; CWBIO, Beijing, China) containing protease inhibitor cocktail tablets (04693132001; Roche, Prague, Czech Republic). Protein concentrations were determined using a BCA assay kit (23225; Thermo Scientific, Waltham, MA, USA). The following antibodies were used for Western blot analysis: anti-MCM3 (#4012; Cell Signaling Technology, Danvers, MA, USA; 1:1000), anti-CDK1 (#9116; Cell Signaling Technology; 1:1000), anti-HIF-1α (#36169; Cell Signaling Technology; 1:1000), anti-phospho-MCM3 (Ser112) (#12686; Cell Signaling Technology; 1:1000), anti-IGF2BP2 (ab124930; Abcam, Cambridge, MA, USA; 1:1000), anti-IGF2BP1 (100999; Abcam; 1:1000), anti-IGF2BP3 (ab177477; Abcam; 1:1000), anti-FTO (712913; Thermo Scientific; 1:1000), anti-MCM2 (A1056; ABclonal, Wuhan, Hubei, China; 1:1000), anti-MCM4 (A13513; ABclonal; 1:1000), anti-MCM5 (A5556; ABclonal; 1:2000), anti-MCM6 (A1955; ABclonal; 1:1000), anti-MCM7 (A1138; ABclonal; 1:1000), anti-H3 (ab1791; Abcam; 1:1000), anti-Lamin B1 (A1910; ABclonal; 1:2000), and anti-β-actin (A5316; Sigma–Aldrich; 1:4000).

To obtain purified MCM3 and HIF-1α proteins, the pGEX-6P-1-MCM3, pGEX-6P-1-HIF-1α or pGEX-6P-1 plasmids were transformed separately into *Escherichia coli* BL21 (DE3) cells. Protein expression was induced with 0.5 mM isopropyl-β-d-thiogalactoside (IPTG) at 25 °C overnight. Finally, proteins were purified on GST purification columns (16107; Thermo Scientific) according to the manufacturer’s instructions. Purified proteins were concentrated with Amicon Ultra0.5 Centrifugal Filter Units (UFC503008; Millipore, Billerica, MA, USA). To explore the direct protein‒RNA interactions in vitro, the purified MCM3 protein was first incubated with a magnetic bead-conjugated antibody. After incubation, the protein sample was divided into two aliquots, and then 2 μg of the in vitro-transcribed LINK-A sense and antisense strands were added separately. The same amount of purified HIF-1α protein was added at the same time. After incubation for 12 hours at 4 °C, the immunoprecipitated protein complexes were washed with Tris-buffered saline with 0.5% Triton X-100 4 times and analyzed by Western blotting.

For the immunoprecipitation (IP) assay, an anti-MCM3 antibody (ab272877; Abcam; 1:100) was incubated with protein A/G magnetic beads (88803; Thermo Scientific) at 4 °C for 6-8 hours. Cells were lysed in IP lysis buffer (P0013; Beyotime Biotechnology, Shanghai, China) containing protease inhibitor cocktail tablets (04693132001; Roche). Equal amounts of lysates were incubated with magnetic bead-conjugated antibodies overnight at 4 °C. Then, the immunoprecipitated proteins were washed with Tris-buffered saline with 0.5% Triton X-100 4 times and analyzed by Western blotting.

RNA preparation and RT‒qPCR were performed according to previously described methods.^27^ The sequences of the primers used for RT–qPCR are listed in Supplementary Table S2.

### m^6^A dot-blot assay

The RNA sample was diluted in the same concentrate in RNase-free water and then denatured at 95 °C for 3 min. Two microliters of RNA was dropped directly onto a Hybond-N+ membrane (RPN119B; GE Healthcare Life Science, Uppsala, Sweden). The RNA samples were cross-linked onto the membrane via 37 °C 30 min dehydration.

The membrane was washed in TBST twice and blocked with 5% nonfat milk for 1 hour at room temperature. The membrane was incubated with m6A antibody (202003; Synaptic Systems, Goettingen, Germany; 1:500) at 4 °C overnight. After incubation, the membrane was incubated with HRP-conjugated goat anti-rabbit IgG and developed with ECL Western Blotting solution. Finally, the membrane was stained with 0.02% methylene blue as a loading control.

### Chromatin-binding protein isolation assay

Chromatin-binding proteins were isolated from cultured cells using a subcellular protein isolation kit (78840; Thermo Scientific) according to the manufacturer’s instructions. In brief, two million ESCC cells were collected, and proteins in the different cell fractions were obtained by successively adding lysates of the different cell fractions and performing differential centrifugation. The final protein lysate of the chromatin-binding component was obtained, and the protein concentration was determined with a BCA kit.

### RNA pulldown and RIP assays

RNA pulldown and RIP assays were performed as previously reported.^27^ Negative control IgG (#2729; Cell Signaling Technology) and anti-MCM3 (ab272877; Abcam; 1:100), anti-IGF2BP2 (ab128175; Abcam; 1:1000) and anti-m^6^A (202003; Synaptic Systems; 1:100) antibodies were used in the RIP assay. MS analysis of proteins precipitated by the LINK-A sense and antisense strands was performed by Shanghai Applied Protein Technology Co., Ltd (Shanghai, China).

### mRNA stability assay

Cells were treated with 5 μg/ml actinomycin D (M4881; Abmole, Houston, TX, USA) to block de novo RNA synthesis. Total RNA was collected at different time points, and the expression of LINK-A was measured by RT‒qPCR. The half-life of RNA was determined by comparing the RNA level at each time point with the RNA level at time 0 normalized to the GAPDH RNA level.

### Detection of RNA synthesis rate

Cell samples were labeled with 0.25 mM EU supplemented growth media for either 4 hours or 5 hours, and the relative RNA synthesis was calculated for 5 hours over a 4-hours pulse duration. After incubation, 90% ethanol was added to fix the cells, and then 0.5% Triton X-100 was used to permeabilize the cells. The cells were treated with the click-reaction solution (0.25 mM biotin-azide, 0.3 mM CuSO_4_, 0.6 mM THPTA, 1 mM aminoguanidine, and 5 mM sodium L-ascorbate). Cells were lysed for 30 min on ice by lysis buffer (20 mM Tris-HCl with pH 7.5, 500 mM LiCl, 1 mM EDTA with pH 8.0, 0.5% lithium-dodecylsulfate (LiDS), and 5 mM DTT). Then, 100 μl of streptavidin-conjugated magnetic beads was added to each dish and incubated for 2 hours at 4 °C, and the cells were washed with lysis buffer 6 times. Finally, RNA elution buffer (10 mM EDTA with pH 8.2 and 95% formamide) was added and incubated at 90 °C for 5 min. RNA extraction and RT‒qPCR were performed according to the above method.

### Dual-luciferase reporter assay

To detect transcriptional activation of hypoxia response elements, the HRE-luciferase plasmid (#26731; Addgene) was transfected into cells with LINK-A-interference. After 48 hours, luciferase activity was measured using a Dual-Luciferase Reporter Assay System (E1960; Promega, Madison, WI, USA). Luciferase activity was calculated according to the ratio of firefly luciferase activity to Renilla luciferase activity.

### FISH and double IF staining

Cells were seeded into a μ-Slide VI (80666; ibidi, Martinsried, Germany) at 60% confluence. After adherence, the cells were fixed with 4% paraformaldehyde and permeabilized with 0.5% Triton X-100 prior to blocking with 5% bovine serum albumin. The FISH assay was performed using a lncRNA FISH Kit (C10910; RiboBio, Guangzhou, Guangdong, China). Hybridization was carried out overnight in a humidified chamber at 37 °C in the dark. After probe binding, the cells were incubated with primary antibodies overnight at 4 °C. Proteins were visualized by incubation with Alexa Fluor 488-conjugated anti-rabbit IgG (#4412; Cell Signaling Technology) or Alexa Fluor 594-conjugated anti-rabbit IgG (#8890; Cell Signaling Technology), and nuclei were stained with Hoechst 33342 (H21492; Thermo Scientific) for 15 min at room temperature. Finally, images were acquired by confocal microscopy.

### Seahorse assay

The OXPHOS and glycolytic capacities were measured with a Seahorse XF96 Extracellular Flux Analyzer (Agilent Technologies, Santa Clara, CA, USA). A Cell Mito Stress Test Kit (103015; Agilent Technologies) and Glycolysis Stress Test Kit (103020; Agilent Technologies) were used to measure the oxygen consumption rate (OCR) and extracellular acidification rate (ECAR), respectively. Specifically, 1×10^4^ KYSE410 or 1.2×10^4^ KYSE450 cells were seeded into Seahorse XF96 Cell Culture Microplates (101085; Agilent Technologies). The ECAR was measured as follows.

After cell adherence, the cell medium was replaced with XF RPMI Base Medium (103575; Agilent Technologies) containing 2 mM glutamate. After calibration was completed, glucose (final concentration: 10 μM), oligomycin (final concentration: 1 μM) and 2-deoxy-D-glucose (2-DG; final concentration: 50 μM) were added successively. The OCR was measured as follows. After cell adherence, the cell medium was replaced with XF RPMI Base Medium (103575; Agilent Technologies) containing 2 mM glutamate, 1 mM sodium pyruvate and 10 mM glucose. After calibration was completed, oligomycin (final concentration: 1.5 μM), carbonyl cyanide 4-(trifluoromethoxy) phenylhydrazone (FCCP) (final concentration: 1 μM), and rotenone/antimycin A (Rot/AA; final concentration: 0.5 μM) were added successively. Wave software (Agilent Technologies) was used to analyze the raw data, and each group was analyzed in biological triplicate.

### RNA-seq

The RNA-seq experiment and data analysis were performed by Shanghai Applied Protein Technology Co., Ltd. Three replicates were prepared for each group using independent cell cultures. The paired-end libraries were prepared using an NEBNext Ultra RNA Library Prep Kit for Illumina (New England Biolabs, Ipswich, MA, USA) following the manufacturer’s instructions, and sequencing was performed using the Illumina NovaSeq 6000 platform. A fold change of >2 and P value of <0.05 were set as the criteria for identifying significantly differentially expressed genes.

### Animal experiments

All animal protocols were approved by the Animal Care and Use Committee of the Chinese Academy of Medical Sciences Cancer Hospital. Cell-derived xenografts and patient-derived xenografts (PDXs) were established as previously described.^46^ Mice bearing PDXs were injected intratumorally twice at 3-day intervals with AAV expressing shCtrl or AAV expressing shLINK-A (2.5 × 10^11^ vg/mouse) produced by ViGene Biosciences (Jinan, Shandong, China). The tumor volume was calculated as 0.52×length×width^2^. The combination index (CI) was calculated as CI=(E1/E2)/(C1/C2), where E1 is the mean tumor volume or weight in the experimental arm (e.g., gene overexpression or knockdown) with drug treatment, E2 is the mean tumor volume or weight in the experimental arm with vehicle treatment, C1 is the mean tumor volume or weight in the control arm (e.g., empty vector or control shRNA) with drug treatment, and C2 is the mean tumor volume or weight in the control arm with vehicle treatment. CI < 1, = 1, and > 1 indicate synergistic, additive, and antagonistic effects, respectively.

### Statistical analysis

Statistical analysis was performed using GraphPad Prism 8.3.0 (GraphPad Software, La Jolla, CA, USA). Unpaired or paired Student’s *t* test, two-way ANOVA, and the Kaplan–Meier method were used for comparisons between two groups, comparisons among multiple groups, and survival analysis, respectively. Statistical significance was assumed for P values ≤ 0.05, and results from at least three biologically independent experiments with similar results are reported. The data are presented as the mean ± s.e.m. or mean ± s.d. values.

## Data availability

The authors declare that the data supporting the findings of this study are available within the main text and supplementary materials.

## Acknowledgments

We are grateful to Dr. Yutaka Shimada (Kyoto University, Kyoto, Japan) for the ESCC cell lines. We thank Dr. Yin Li (Cancer Hospital, Chinese Academy of Medical Sciences, Beijing, China) for the ESCC samples. We appreciate Shanghai Applied Protein Technology Co., Ltd. (Shanghai, China) for performing the MS analysis and RNA-seq. Q.L. is supported by a Fellow grant from the Leukemia & Lymphoma Society. This study was supported by funding from the National Key R&D Program of China (2021YFC2501000), the National Natural Science Foundation of China (82030089, 82188102, 82172590), the CAMS Innovation Fund for Medical Sciences (2021-I2 M-1-018, 2021-I2 M-1-067), and the Fundamental Research Funds for the Central Universities (3332022135).

## Author Contributions

**Y.N**., **S.L**., **Q.L**. and **X.W**. designed the experiments and analyzed the data. **Y.N**. and **S.L**. performed most of the experiments. **W.C**. and **P.Z**. assisted with the animal studies. **Y.N**. drafted the manuscript with assistance from **S.L**. and **Z.L**. All authors reviewed the manuscript. **Z.L**. organized and supervised this study.

## Declaration of Interests

The authors declare no conflicts of interest.

## Notes

### Competing Interest Statement

The authors have declared no competing interest.

