## Supplemental Figures and Tables for "m^6^A Demethylase FTO Stabilizes LINK-A to Exert Oncogenic Roles via MCM3-Mediated Cell Cycle Progression and HIF-1α Activation"

### Supplementary figures

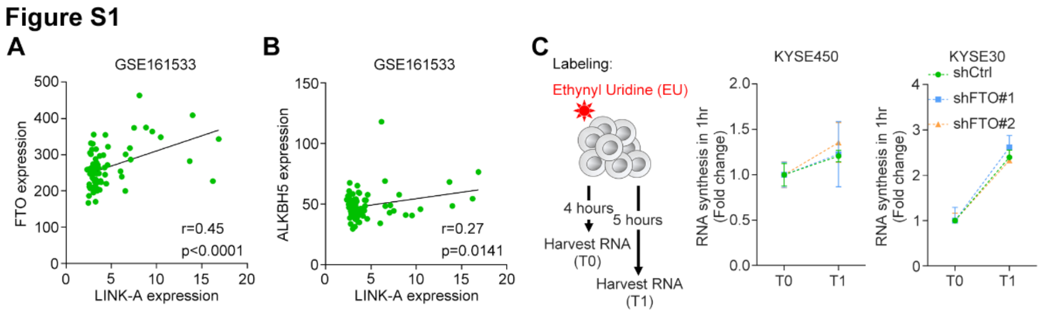


**Figure S1. FTO stabilizes LINK-A by m6A demethylation.** (A and B) Correlations between LINK-A expression and FTO or ALKBH5 expression were determined using Pearson correlation analysis in the GEO database. (C) Dynamics of RNA synthesis calculated for the indicated cells. Cartoon of the experimental scheme is shown on the left. The data are presented as the mean ± s.d. values; n = 3.

**Figure S2**

**A**

| Variable | | LINK-A (± s.e.m.) | p value |
|---|---|---|---|
| Grade | Well/Moderate | 25.22±2.022, n=158 | 0.2122 |
| | Poor | 30.64±4.487, n=55 | |
| Tumor depth | T1/T2 | 25.00±3.535, n=59 | 0.9590 |
| | T3/T4 | 25.21±2.091, n=166 | |
| Lymph node metastasis | N0 | 22.03±2.812, n=96 | 0.0189 |
| | N1 | 31.58±2.891, n=95 | |
| Distant metastasis | M0 | 27.20±2.087, n=184 | 0.2914 |
| | M1-2 | 15.71±9.157, n=7 | |

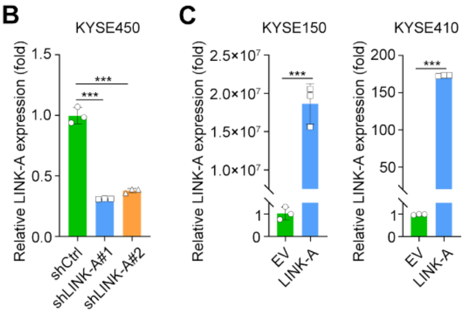


**Figure S2. LINK-A promotes ESCC malignancy.** (A) Correlation analyses of LINK-A expression with other clinicopathological features in patients with ESCC. Unpaired Student's *t* test. (B and C) RT–qPCR analysis of LINK-A expression in indicatted cells. The data are presented as the mean ± s.d. values; unpaired Student's *t* test, ***$p < 0.001$; $n = 3$.

**Figure S3**

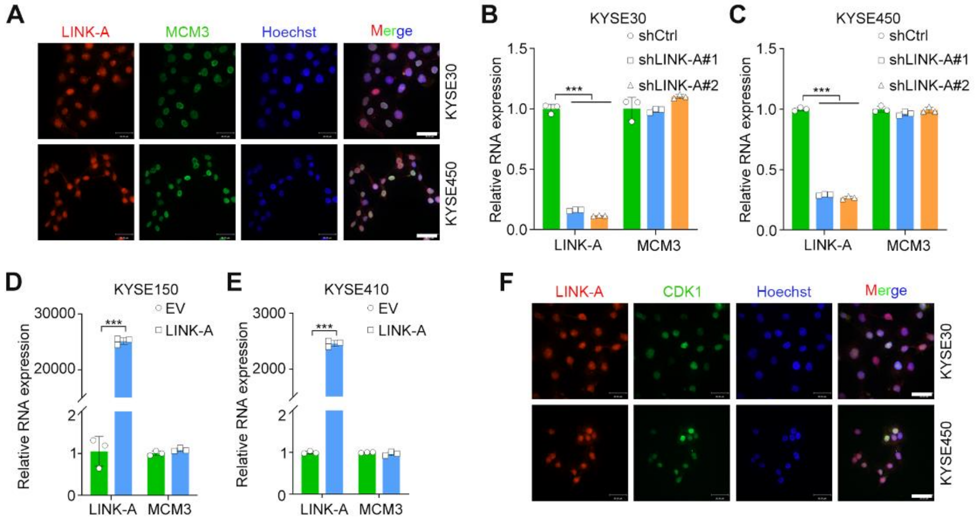


**Figure S3. LINK-A promotes MCM3 phosphorylation by mediating the interaction between MCM3 and CDK1.** (A) IF analysis of LINK-A (red) and MCM3 (green) in the indicated cells. Nuclei (blue) were labeled with Hoechst. The scale bar represents 30 μm. (B-E) RT–qPCR analysis of LINK-A and MCM3 expression in the indicated cells. The data are presented as the mean ± s.d. values; unpaired Student's *t* test, ***p < 0.001; n = 3. (F) IF analysis of LINK-A (red) and CDK1 (green) in the indicated cells. Nuclei (blue) were labeled with Hoechst. The scale bar represents 30 μm.

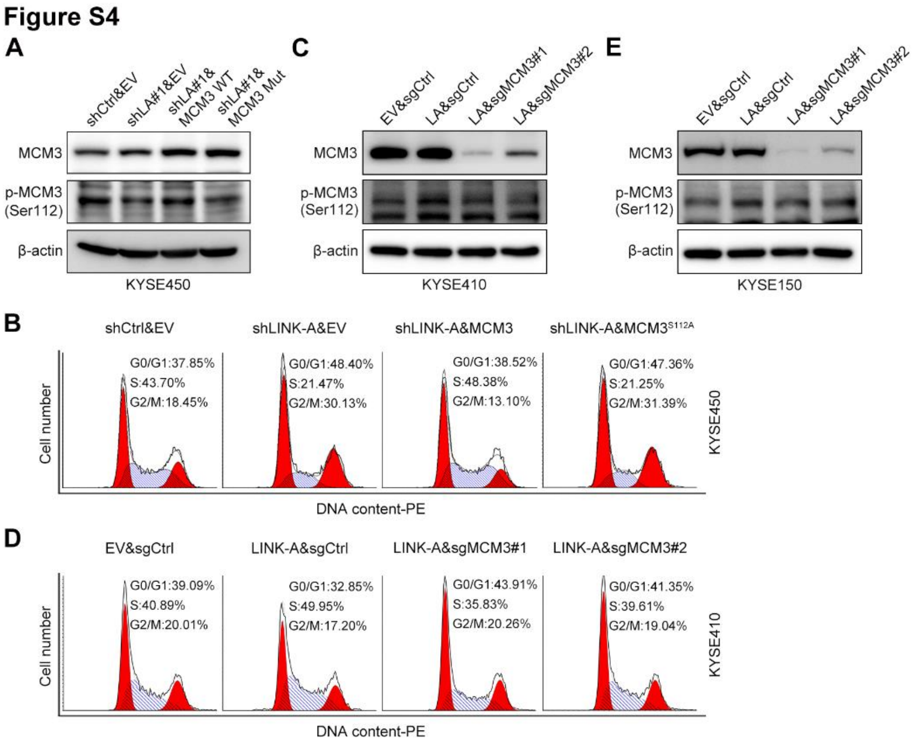


**Figure S4. MCM3 mediates LINK-A to facilitate cell cycle progression.** (A, C and E) Western blot analysis of MCM3 and phosphorylated MCM3 in the indicated cells. (B and D) Representative images of cell cycle analyses in the indicated cells.

**Figure S5**

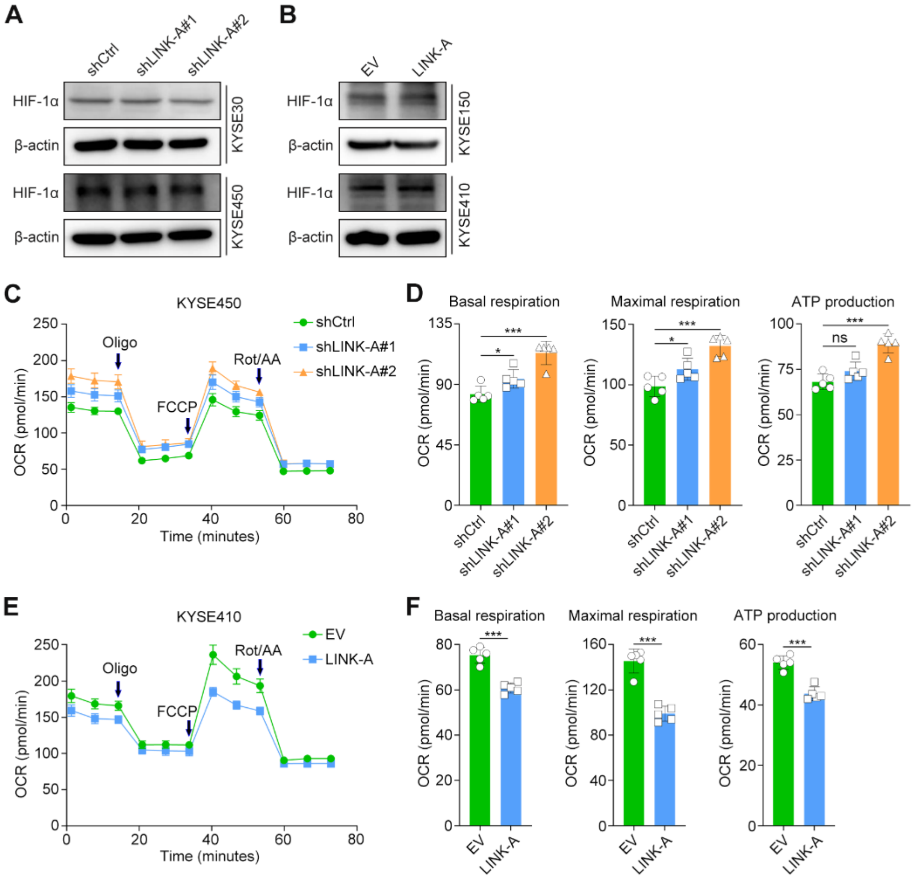


**Figure S5. LINK-A induces HIF-1α transcriptional activity by sequestering HIF-1α from MCM3.** (A and B) Western blot analysis of HIF-1α expression in the indicated cells. (C-F) Measurement of the OCR and mitochondrial respiration level using a Seahorse assay in LINK-A-silenced KYSE450 cells (C and D) and LINK-A-overexpressing KYSE410 cells (E and F). The data are presented as the mean ± s.d. values; unpaired Student's *t* test, * $p < 0.05$, ***$p < 0.001$, ns: no significance; n = 5.

### Supplementary tables

#### Table S1. shRNA and sgRNA sequences

| Sequence (5' to 3') | Name |
|---|---|
| AGATGTAGTTCTAGTTCAT | shLINK-A#1 |
| GGTCTTCATTCTTACGCTT | shLINK-A#2 |
| CCCATTAGGTGCCCATATTTA | shFTO#1 |
| CGGTTCACAACCTCGGTTTAG | shFTO#2 |
| GCTGGACGATGTGGAGCTGC | sgMCM3#1 |
| GCACCGTTACAGAGCACCTG | sgMCM3#2 |

**Table S2. RT–qPCR primers**

| Sequence (5' to 3') | Name |
|---|---|
| CCGGGAAACTGTGGCGTGATGG | GAPDH-F |
| AGGTGGAGGAGTGGGTGTCGCTGTT | GAPDH-R |
| CTCGCTTCGGCAGCACA | U6-F |
| AACGCTTCACGAATTTGCGT | U6-R |
| CCTCTGACTGATGCTTCC | LINK-A-F |
| CACCTGGCTGTCTTATTCC | LINK-A-R |
| AGTTCGTCCCAAAGTCGTCC | MCM3-F |
| GGGGATTGTTCTCCTCATCCT | MCM3-R |
| AGAATGTCTGTGACGATGTGG | FTO-F |
| GCACTTTCTGTATCGATTGCC | FTO-R |
| CGGCGAAGGCTACACTTACG | ALKBH5-F |
| CCACCAGCTTTTGGATCACCA | ALKBH5-R |
| TGCTTGCCAAAAGAGGTGGA | HIF-1α-F |
| TTCTGTGTCGTTGCTGCCAA | HIF-1α-R |
| ACCAGGACAGCCAATACAAG | PDK1-F |
| CCTCGGTCACTCATCTTCAC | PDK1-R |
| ATCTTGACCTACGTGGCTTGGA | LDHA-F |
| CCATACAGGCACACTGGAATCTC | LDHA-R |
| CGCAAGAAATCCCGGTATAA | VEGF-F |
| TCTCCGCTCTGAGCAAGG | VEGF-R |

| | |
|---|---|
| GGTGGGGTCATGTGTGTGG | BCL2-F |
| CGGTTCAGGTACTCAGTCATCC | BCL2-R |
| TGAGCATCGTGGCCATCTTT | SLC2A1-F |
| CCGGAAGCGATCTCATCGAA | SLC2A1-R |
| TGCCACCAGACTAAACTAGACG | HK2-F |
| CCCGTGCCCACAATGAGAC | HK2-R |
| CCAGGGATTTCAGTCGATGT | Firefly-Luc-F |
| AATCTCACGCAGGCAGTTCT | Firefly-Luc-R |
| GTGGTGGGCCAGATGTAAAC | Renilla-Luc-F |
| ACCAGATTTGCCTGATTTGC | Renilla-Luc-R |
